# Comprehensive analysis of yeast +1 ribosomal frameshifting unveils a novel stimulator supporting two distinct frameshifting mechanisms

**DOI:** 10.1101/2025.03.19.644219

**Authors:** Darren A. Fenton, Maria Bożko, Michał I. Świrski, Gary Loughran, Martina M Yordanova, Joanna Kufel, John F Atkins, Pavel V. Baranov

## Abstract

Ribosomal frameshifting is an important, albeit rare, mRNA decoding mechanism that generally allows the synthesis of a single protein from two different reading frames. For most +1 frameshifting cases, the mechanism is commonly presumed to involve dissociation of the P-site tRNA from its cognate codon followed by its movement to the +1 codon, setting the new +1 frame for incoming tRNAs. This movement is stabilized by P-site tRNA pairing with the +1 codon. However, in several occurrences in the yeast *Saccharomyces cerevisiae*, P-site tRNA re-pairing with the +1 codon is impossible. Two alternative hypotheses exist explaining this observation. One model suggests that +1 frameshifting occurs according to a common mechanism involving P-site movement, while its re-pairing with +1 codon is not essential. The alternative model suggests a distinct mechanism in which the A-site tRNA acceptance at the +1 codon occurs in the absence of P-site tRNA movement relative to mRNA. Here we set out to perform a comprehensive comparative analysis of all known +1 ribosomal frameshifting sites in *S. cerevisiae*. This included a novel case of +1 ribosomal frameshifting that we discovered during this study. It is required for the expression of *LLP1* gene encoding dolichol-linked oligosaccharide pyrophosphatase. During the analysis of all frameshifting contexts, we identified a conserved RNA secondary structure located almost immediately upstream of the ABP140 frameshifting site. This structure substantially increases +1 frameshifting efficiency. The RNA stimulator’s location suggests that mRNA exiting the ribosome forms this structure, creating an mRNA pulling effect, thus favouring positioning of the +1 codon in the P-site. Placing the stimulator upstream of various known frameshifting sites, revealed that its stimulatory action is selective to those frameshifting sites where P-site tRNA re-pairing is possible, reinforcing the idea of two distinct mechanisms of ribosomal frameshifting.

## Introduction

Translating ribosomes are capable of deviating from the “standard” decoding rules in response to specific mRNA signals. These deviations are collectively termed “recoding events” (1–3). Ribosomal frameshifting is a form of recoding where some ribosomes shift reading frame at a specific site during translation, resulting in the creation of a trans-frame protein in addition to the standard translation product. While ribosomal frameshifting is common in mobile elements and especially in RNA viruses (4), it is extremely rare in chromosomal gene decoding, with the exception of ciliates *Euplotes* where it is considered a feature of its genetic code (5). Normally, frameshifting occurs at a specific combination of (i) codons that allow tRNA repositioning with codons in a new frame and (ii) stimulatory signals that elevate its efficiency through a variety of molecular mechanisms.

The first example of a eukaryotic gene requiring frameshifting was discovered in the *Saccharomyces cerevisiae* transposable element Ty1, which utilizes efficient +1 ribosomal frameshifting (6–9). Strikingly, a minimal sequence of 7 nucleotides was found to be sufficient to support a remarkably high (∼40%) frameshifting efficiency in the absence of additional RNA stimulators in the vicinity of the frameshifting site (10). The sequence of the Ty1 frameshift heptamer is CUU_A.GG_C where underscore indicates codon boundaries in the initial (0) reading frame and dot indicates codon boundaries in the new (+1) frame (this notation will be used thereafter). A remarkable feature of this frameshifting site is the high imbalance in the levels of tRNAs recognising the AGG (Arg) and GGC (Gly) codons. The *S. cerevisiae* genome contains only a single copy of Arg-tRNA (CCU) gene, while the number of gene copies of Gly-tRNA (GCC) is up to 18 depending on the strain (11). This suggests a far greater concentration of tRNAs cognate for GGC over tRNAs cognate for AGG and subsequently faster decoding of the GGC codon in comparison with the AGG. Presumably, slow decoding of AGG allows more time for re-pairing of the Leu-tRNA in the P-site from CUU to UUA while fast decoding of GGC would prevent re-pairing in the reverse direction from UUA to CUU (10,12). Subsequently, the same frameshift-inducing heptamer was identified in the genes *ABP140* (13) and *YPL034W*/*YFS1* (14). Here, we also report its utilization for the expression of *YJR112W-A*/*LLP1*. A different pattern, using the same P-site codon but different A-site codons (CUU_A.GU_U) was found to cause +1 frameshifting in *EST3* (15). Albeit different and less efficient, the mechanism of *EST3* frameshifting may be similar. In *S. cerevisiae*, AGU decoding requires wobble interactions and may be slow, while the number of copies for the cognate Val-tRNA is high, reaching up to 21 (11).

A more different +1 frameshifting heptamer (GCG_A.GU_U) was identified in the Ty3 transposable element (16). While the A-site component of this pattern is identical to *EST3* frameshifting, the P-site codon (GCG) is different and seemingly does not support re-pairing in the +1 reading frame. Importantly, while the optimal Ala-tRNA (UGC) for GCG is present, it is rare. The same codon can be decoded by a near cognate isoacceptor tRNA with anticodon IGC which is apparently responsible for frameshifting since it is inhibited by the oversupply of Ala-tRNA (UGC) (17). The same P-site codon is used in the *S. cerevisiae* antizyme (*OAZ1*/*YPL052W*) frameshifting heptamer (GCG_U.GA_C) where frameshifting is driven by competition between termination at a stop codon in poor context and incorporation of a tRNA at the +1 codon. The efficiency of antizyme frameshifting is sensitive to polyamine levels (18–20). The discovery of an unusual P-site codon in the Ty3 frameshifting heptamer prompted a subsequent investigation of the repertoire of P-site codons supporting +1 ribosomal frameshifting. It revealed several P-site codons supporting above-background levels of +1 frameshifting in A.GU_U or A.GG_C A-site contexts with little P-site re-pairing potential with the +1 codon (21). This led to the suggestion that certain tRNAs, when in the P site, possess properties that allow ribosomes to incorporate A-site tRNAs in the +1 frame without prior P-site tRNA slippage (22). To unify these observations into a single parsimonious mechanism, kinetic considerations have been used to argue that it is the dissociation of the P-site tRNA from its zero-frame codon that is the limiting step. Even short-lived unstable P-site tRNA interactions with the +1 codon may be sufficient for high efficiency frameshifting given the high imbalance between cognate in- and out-of-frame tRNAs in the A-site (12). However, until now, little experimental evidence has been provided to resolve this conundrum. It has remained unclear whether one or two different mechanisms are responsible for +1 ribosomal frameshifting in yeast at different P-site codons (Fig. 1).

**Figure 1.**
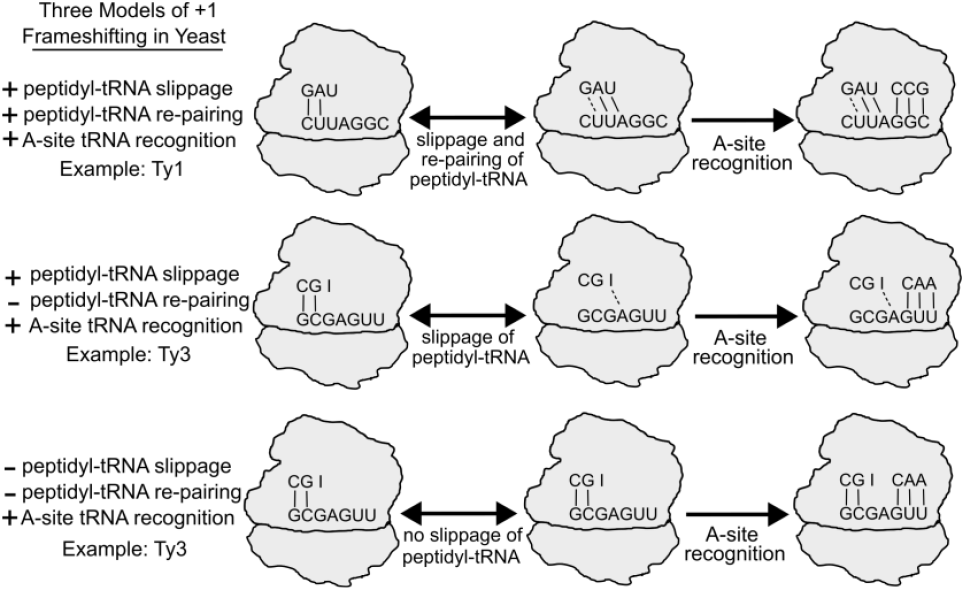
Models of +1 Frameshifting Mechanisms in *Saccharomyces cerevisiae*. Top model: Peptidyl-tRNA slippage and re-pairing to the overlapping UUA codon in the +1 frame is followed by A-site recognition of the tRNA decoding the +1-frame GGC codon. The middle and bottom schematics depict two alternative mechanisms proposed to explain frameshifting when tRNA re-pairing with the +1 codon in the P-site could not occur. Middle model: Peptidyl-tRNA slippage, with no re-pairing, followed by A-site recognition of the tRNA decoding the +1-frame GUU codon. Bottom model: +1 frameshifting involves no slippage or re-pairing of the peptidyl-tRNA. Instead, frameshifting occurs due to direct recognition of tRNA to the +1 codon in the A-site.

In this study, our initial motivation was to carry out a systematic comparative study of naturally occurring +1 frameshifting sites, with a goal to provide a reliable reference for +1 frameshifting efficiencies in yeast. This was due, since prior frameshifting measurements were made in different research groups using different reporter systems that may be subject to various technical artifacts (23), and thus could not be directly compared. For this purpose, we have developed a new vector designed for studying recoding signals in yeast, based on the StopGo reporter system previously developed for mammalian cell lines (24). Using this reporter system, we measured ribosomal frameshifting at all the above-mentioned cases with and without their surrounding mRNA context. The exploration of the contexts revealed the existence of a previously unknown +1 frameshifting stimulator within the *ABP140* mRNA, located upstream of the frameshifting site heptamer. This stimulator increases +1 frameshifting efficiency from 40% to 60%. Phylogenetic analysis of the corresponding sequence combined with site-directed mutagenesis strongly suggest that this stimulator is an RNA structure whose formation pushes the ribosome downstream into the +1 frame. We tested the effect of this stimulator on different frameshifting heptamers allowing us to discriminate frameshifting occurring due to P-site tRNA slippage or direct out-of-frame A-site tRNA recognition.

In addition to resolving a long-standing mechanistic conundrum, our study provides a reference to all *S. cerevisiae* +1 frameshifting cases reported so far, including a novel case we discovered while exploring available ribosome profiling data. Finally, ribosome profiling data was used as an orthogonal approach for estimating endogenous levels of ribosomal frameshifting.

## Methods

### Cloning

SnapGene (www.snapgene.com) was used to codon optimise (based on codon usage) Renilla and Firefly luciferase sequences from pSGDLuc (24) and synthesized as a gblock from IDT (Integrated DNA Technologies, Leuven, Belgium). The reporter cassette was inserted into the pGREG-505 plasmid which was linearized with SalI (NEB, #R3138S) and size selected to remove the *HIS3* coding sequence. For insertion of short frameshifting heptamers, annealed oligos were ligated into the NdeI (NEB, #R0111S) and NheI (NEB, #R3131S) digested vector with T4 DNA ligase (NEB, #M0202S) and incubated overnight at room temperature in a 5 μL reaction. For insertion of longer frameshifting cassettes, 60 ng of NdeI and NheI digested plasmid and ∼7 ng of ∼200 bp inserts were added to 15 μL of homemade Gibson assembly master mix (25) and incubated at 50°C for 1 hour in a thermocycler. 5 μL of the Gibson assembly or T4 ligation reaction was transformed into chemically competent *E. coli* DH5a cells. The empty pYSGDLuc plasmid is available from Addgene (Plasmid #236297).

### Yeast culturing and transformation

5 mL of YPD broth was inoculated with a single colony of *S. cerevisiae* (strain BY4741) and incubated overnight at 30°C and 200 RPM. The next day, 200-400 ng of plasmid was transformed into cells *via* the Lithium acetate/Salmon sperm carrier method (26). Cultures were plated onto SC 2% glucose media with an amino acid drop out -Leucine mix (Formedium, #DSCK052) agar plates and incubated at 30°C for 2-3 days until colonies formed.

### Luciferase Assays

Two independent yeast colonies were inoculated in 5 mL 2% glucose -Leucine medium in a 50 mL Falcon tube and incubated overnight at 30°C and 200 rpm. The next day, the A600 (O.D. 600) values were measured and the culture was diluted to an A600 of ∼0.1. For the 96 well plate assay, 200 μL of culture was plated into each well (a total of six wells per biological replicate). The plate was sealed using gas-permeable tissue culture seals (4titude, #4ti-0516/384), placed on a plate shaker inside a 30°C incubator and incubated for ∼6 hours. 50 μL of the culture was then transferred to a white full area 96 well plate and 50 μL of 2X Passive Lysis Buffer (Promega) was added to each well and incubated for 50-60 minutes with shaking at room temperature to allow lysis. 50 μL of homemade LAR (substrate for Firefly luciferase) and 50 μL homemade StopGlow (substrate for Renilla luciferase) were injected into each well (27).

### Western Blotting

Yeast transformants were inoculated in 5 mL -Leucine media in a 50 mL falcon tube and incubated overnight at 30°C with 200 RPM. A600 values were measured and ∼10 A600 units were transferred to a 1.5 mL tube. Cells were washed once in sterile water and boiled in 280 μL of 1X SDS-sample buffer for 3 minutes. 5-10 μL was transferred onto NuPage protein gels (Invitrogen) and run for 200V for ∼24 minutes with 1X MES running buffer (Invitrogen). Proteins were transferred to a Nitrocellulose membrane using a BioRad Transblot. Membranes were blocked with 5% low fat milk in 1X PBS-T for 45 minutes at room temperature with gentle agitation. The membranes were treated with a primary antibody solution consisting of mouse anti-Renilla (Millipore) and goat anti-Firefly (Promega) in 1% BSA and PBS-T and was incubated overnight at 4°C. The membrane was washed three times in PBS-T for 5 minutes each before and after secondary antibodies (anti-mouse red and anti-goat green). Membranes were visualized using an Odyssey imager. Densitometry analyses were carried out using imageJ.

### Ribosome Profiling Analysis

Data from each study was obtained via the Ribocrypt browser (https://ribocrypt.org/). To determine the +1-frameshifting efficiency, the number of RPFs on each frame was normalized by the number of codons. The normalized number of RPFs in the +1 frame was divided by the number in the 0-frame and multiplied by 100 to obtain the % frameshifting.

### Bioinformatic and Computational Analyses

ABP140 homologs were found using TBLASTN (28) against the genomes of budding yeast species. To produce in-frame alignments for homologs that do not contain frameshifting, the +1-frame was fused to the 0-frame by replacing CTTAGGC with CTTGGC. RNAfold was used to determine RNA secondary structures (29). Orthologous sequences were aligned using MUSCLE (30) and converted to a codon alignment using PAL2NAL (31). A codon alignment viewer was generated with python. The rate of synonymous substitutions across multiple sequence alignment was analysed with Synplot2 (32), using a window size of 17 codons. DMS-seq data was obtained from (33) and the Varna software was used to visualize RNA structure and DMS-seq data (34).

## Results

### A new reporter to test recoding efficiencies in *S. cerevisiae*

Measurements of frameshifting efficiencies are usually obtained using vectors that contain candidate frameshifting sequences fused between two expression reporters. Previously used reporters in yeast frameshifting studies included β-galactosidase, luciferase, β-galactosidase-luciferase fusions, and dual-luciferase reporter (7,35,36). Luciferase assays are attractive as they allow sensitive readings, especially for cases where a low recoding signal needs to be measured. An in-frame control (i.e. both reporters in the same frame) is used to establish the “100% efficiency” signal, allowing comparison with out-of-frame reporters to determine frameshifting efficiency. However, the sequence of the chimeric protein encoded by the test sequence could affect the activities of one or both reporters, leading to alterations in measured activities and distorting accurate measurements of frameshifting (23). To mitigate this issue, the original dual luciferase reporter vector (pDLuc) developed for use in mammalian models (37,38) was modified to incorporate StopGo/2A on both sides of the test sequence in pSGDLuc vector (24). The StopGo/2A peptide motif results in the synthesis of two separated protein products (encoded upstream and downstream of it) because the translating ribosome fails to form a peptide bond at a specific position (reviewed in Luke and Ryan, 2024). These motifs were originally found in viruses where they are responsible for the production of distinct products from the same translated ORF (40). Therefore, most reporter proteins produced from the in-frame and test sequence are identical, allowing a more accurate determination of the frameshifting efficiencies. Indeed, the use of StopGo/2A-containing dual luciferase reporter led to identification of a false positive frameshifting case in a human gene due to limitations of the previous reporter (41).

We adapted the mammalian pSGDLuc reporter plasmid for use in *S. cerevisiae* with three modifications. 1). We optimised Renilla and Firefly luciferase coding sequences by introducing synonymous codon substitutions based on yeast codon usage (Methods). 2). As the StopGo/2A motif used in the mammalian reporter (F2A, derived from foot-and-mouth disease virus) is only ∼50% efficient in *S. cerevisiae*, we substituted it with a more efficient variant, E2A (derived from the equine rhinitis A virus) which is ∼90% efficient (42). The new vector backbone encodes both leucine and G418 selection markers (Fig. 2A). 3). We modified a multiple cloning site to incorporate the NheI and NdeI restriction enzyme sites to allow cloning via T4 ligation or Gibson assembly reactions. The new vector termed pYSGDLuc (Fig. 2A) can be used for various types of recoding studies in yeast.

**Figure 2.**
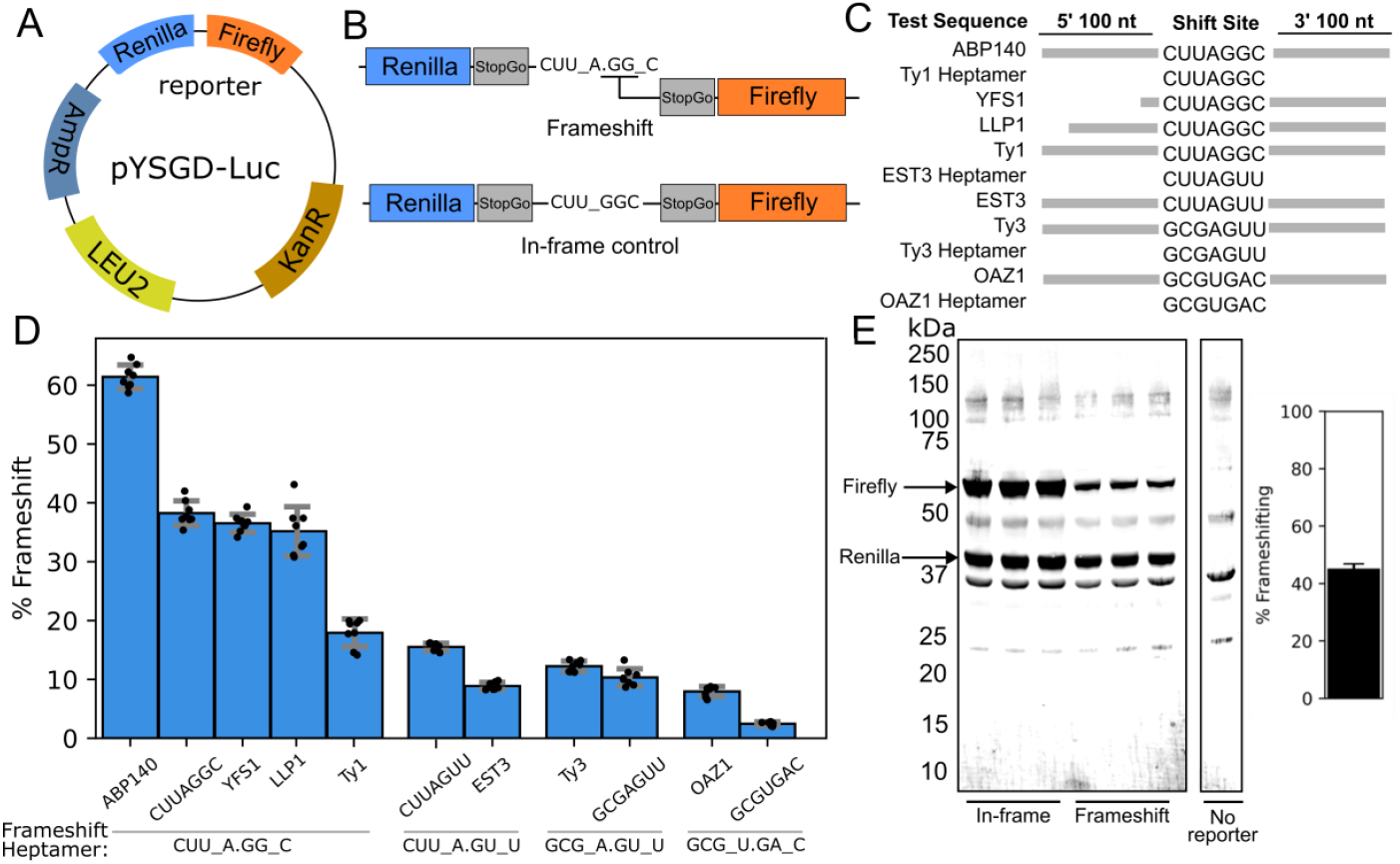
Uniform assessment of frameshifting efficiency at known frameshifting cases. **A**, A schematic representation of the pYSGDLuc. *LEU2* represents the Leucine biosynthesis gene and AmpR and KanR represent ampicillin (for *E. coli* cloning) and kanamycin resistance (for yeast G418 selection) genes. **B**, Schematic of frameshifting reporters for testing the Ty1 heptamer sequence, including WT and mutant in-frame control sequences. **C**, Representation of natural frameshifting cassettes tested in this study. Grey bars indicate the length of tested sequences flanking heptamers. **D**, Frameshifting efficiencies determined with luciferase assays. Cases are clustered together based on the frameshifting heptamer (for example, *ABP140, YFS1, LLP1* and Ty1 frameshifting at CUU_A.GG_C). Error bars represent standard deviation (n=8). **E**, Western Blotting of replicate cell extracts derived from cells expressing the Ty1 heptamer (CUU_A.GG_C) reporters, using anti-Renilla and anti-Firefly antibodies. Densitometry analyses estimate ∼45% frameshifting.

### Comparative analysis of S. cerevisiae frameshifting cassettes using pYSGDLuc and publicly available ribosome profiling data

Using pYSGDLuc, we performed a comparative study of all known naturally occurring cases of efficient +1 frameshifting in *S. cerevisiae*. Measuring these cases in the same reporter system under the same environments (e.g. yeast strain, media, equipment and reagents), allows objective comparison of frameshifting efficiencies, that are not biased by differences in environmental parameters. Each tested case included both an in-frame control and frameshifting test sequence as recommended by recently developed guidelines (23), see Fig. 2B. For all cases, we tested the minimum heptamer sequence (e.g. CUU_A.GG_C) required for frameshifting that includes the P-site and A-site codons before and after frameshifting (Fig. 2C). This provided the basal rate of frameshifting at corresponding heptamers regardless of their natural surrounding nucleotide context. To assess the effect of the frameshifting heptamer within the native mRNA context, we included a ∼100 nt flanking region. The frameshifting site for *YFS1* is located close (∼42 nt) to the 5’ end of the mRNA transcript, therefore, a shorter upstream test sequence was included. The frameshifting efficiencies calculated as a ratio of luciferases ratios between test sequences and controls are shown in Figure 2D as percentages.

To test whether these reporters may produce unexpected proteins, we carried out western blotting to confirm the sizes of the Renilla and Firefly reporter proteins for the TY1 heptamer-only reporter (Fig. 2E, left). Densitometry analyses estimated similar frameshifting efficiencies as the luciferase assays (Fig. 2E, right).

The measured differences between four frameshifting heptamer-only reporters appeared to be consistent with previous reports, CUU_A.GG_C ∼40% > CUU_A.GU_U ∼15% > GCG_A.GU_U ∼10% > GCG_U.GA_C ∼3% (Fig. 2D). Testing frameshifting efficiencies at these heptamers demonstrated a certain degree of context dependence. The inclusion of natural sequence contexts of Ty3, *YFS1* and *LLP1* (Fig. 2C) does not alter basal levels of frameshifting at their corresponding frameshifting heptamers. However, frameshifting in the *ABP140* context was increased 1.5-fold to ∼60% and unsurprisingly, nearly 3-fold for *OAZ1* in its natural context to ∼8%. The Ty1 and *EST3* contexts had opposite effects, with TY1 reducing frameshifting efficiency 2-fold to ∼20% and 1.5-fold to 10% in *EST3*.

Inclusion of only the surrounding 200 nt surrounding sequence into pYSGDLuc cassette does not entirely recreate the parameters of natural frameshifting for at least two reasons. First, sequence elements at large distances from the frameshifting site may alter its efficiencies (43). Second, the efficiency of frameshifting may depend on parameters of mRNA translation not directly linked to the surrounding sequence, such as ribosome loading (44). Therefore, to assess the natural levels of frameshifting we used publicly available ribosome profiling data and inferred frameshifting efficiency from a change in ribosome footprint density downstream of frameshifting sites, similarly to previous work on viral frameshifting (45), see Methods. A global aggregate of these ribosome profiling data from different studies for *ABP140, EST3* and *OAZ1* is presented in Figure 3A. For most cases, we found a strong concordance between frameshifting efficiencies measured with these two methods (Fig. 3B), suggesting that our pYSGD-Luc reporter provides accurate measurements of frameshifting efficiencies and supporting the absence of relevant context outside of the 200 nt proximal window. We note that for *YFS1* and *YJR112W-A/LLP1*, the short length of coding region upstream of frameshifting sites limits the accuracy of this method in comparison with *ABP140, EST3* and *OAZ1*. Nonetheless, for *ABP140, EST3* and *OAZ1*, we find a strong concordance in frameshifting efficiencies between luciferase assays and ribosome profiling.

**Figure 3.**
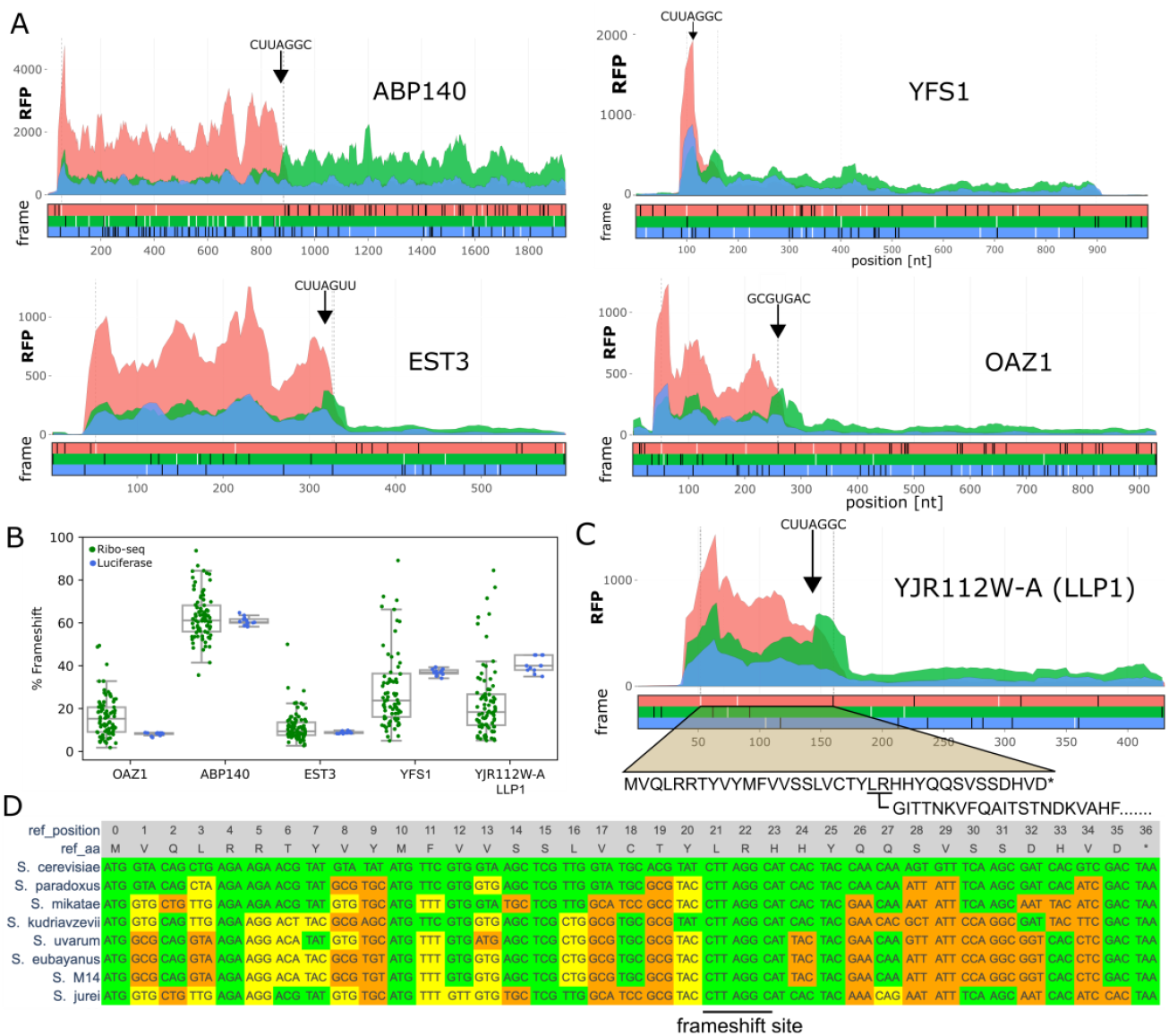
Analysis of publicly available ribosome profiling data. **A.** Subcodon ribosomal profiles for *ABP140, EST3, YFS1* and *OAZ1* genes. For each gene, top plots show aggregated ribosome profiling data differentially coloured based on the supported reading frame. The colours are matched to the reading frames in the ORF plot below where black and white dashes represent stop and ATG codons in each reading frame respectively. The positions of the frameshifting sites are denoted with black arrows, stop codons of the zero frame ORFs are indicated with grey vertical lines. **B**. Ribosomal frameshifting efficiencies inferred from 85 individual ribosome profiling datasets based on the relative ratio of footprint densities in the +1 ORF downstream of the zero frame ORF are shown as green dots. Blue dots indicate frameshifting efficiencies obtained with dual luciferase assays for each replicate (n=8-12). The parameters of distribution for both types of data are shown with boxplots where centre line represents the median, box limits indicate the 25th and 75th percentiles and whiskers extend 1.5 times the interquartile range from the 25th and 75th percentiles. **C**. Same as in A, but for novel instance of frameshifting discovered in this study during translation of the *LLP1* mRNA. Note the full coding regions are not displayed. **D**. Codon alignment of the 0-frame ORF of *LLP1* orthologs in *Saccharomyces* genus with the universally conserved CUU_A.GG_C indicated. Yellow codons represent synonymous changes and orange codons represent non-synonymous changes.

### A novel case of cellular +1 frameshifting at the YJR112W-A/LLP1 locus

Manual analysis of ribosome profiling data revealed an unusual distribution of ribosome-protected fragments (RPFs) in the *YJR112W-A* locus in *S. cerevisiae* (Fig. 3C). *YJR112W-A* is annotated as an intron-containing gene (see Supplementary Fig. S1). However, publicly available RNA-seq data in GWIPS-viz (46) suggest that the annotated intron is a part of mRNA as RNA-seq density is uniform across intronic and exonic regions of this gene (Supplementary Fig. S1). The examination of this ORF sequence revealed the presence of the CUU_A.GG_C heptamer suggesting that ribosomes translating this ORF undergo +1 ribosomal frameshifting. Indeed, the ribosome footprint density is present downstream of this ORF matching the +1 reading frame, albeit at a lower density than at the zero frame ORF. We conclude that the current *S. cerevisiae* reference annotation of *YJR112W-A* CDS is incorrect, and that *YJR112W-A* does not contain introns, instead its CDS is comprised of two overlapping ORFs translated *via* ribosomal frameshifting. The sequence alignments of *YJR112W-A* orthologs from other *Saccharomyces* demonstrated the universal conservation of the CUU_A.GG_C pattern in these species (Fig. 3D) indicating that it evolves under purifying selection suggesting the functional importance of this frameshifting site for these species’ fitness. When tested in its native context, *YJR112W-A* frameshifting is approx. 40% efficient, similar to the efficiency of the CUU_A.GG_C heptamer. Therefore, it is unlikely this novel case contains proximal stimulating or attenuating sequence elements as tested in our luciferase system. While we were preparing a revised version of this manuscript, an independent discovery of +1 frameshifting in *YJR112W-A* locus was reported and the encoded protein was named Llp1 (*LLP1*) (47). Both *YFS1* and *LLP1* contain the CUU_A.GG_C frameshifting heptamer close to the 5’ end of the mRNA transcripts.

### The ABP140 mRNA contains a frameshifting stimulator 5’ of its shift site

Surprisingly, *ABP140* +1 frameshifting is 60% efficient despite containing the same frameshifting heptamer, CUU_A.GG_C, whose frameshifting efficiency is ∼40%, suggesting a stimulatory frameshifting element within the flanking 200 nt native sequence. The hypothesis of a stimulator in ABP140 is further supported by frameshifting efficiencies estimated from ribosome profiling data (Fig. 3A,B). We used RNAfold (29) to predict potential RNA secondary structures within flanking regions and identified a potential stable RNA stem-loop 6 nt upstream of the frameshifting heptamer (Fig. 4A). The existence of the stem-loop is supported biochemically by DMS-seq data, showing a lack of unpaired A and C bases in the stem regions relative to the loop and internal loops (Fig. 4A). An alignment of *ABP140* sequences from multiple *Saccharomyces* species shows conservation of the sequence specifying the predicted stem-loop at this position with several matched substitutions that maintain potential base pairing (Fig. 4D). We used Synplot2 (32), to determine whether there is an increased purifying selection acting on synonymous positions within *ABP140* which would be expected in the presence of functionally significant RNA secondary structure. It reveals a statistically significant decrease in the synonymous substitution rates in comparison with regions outside of the potential RNA secondary structure (Fig. 4B,C).

**Figure 4.**
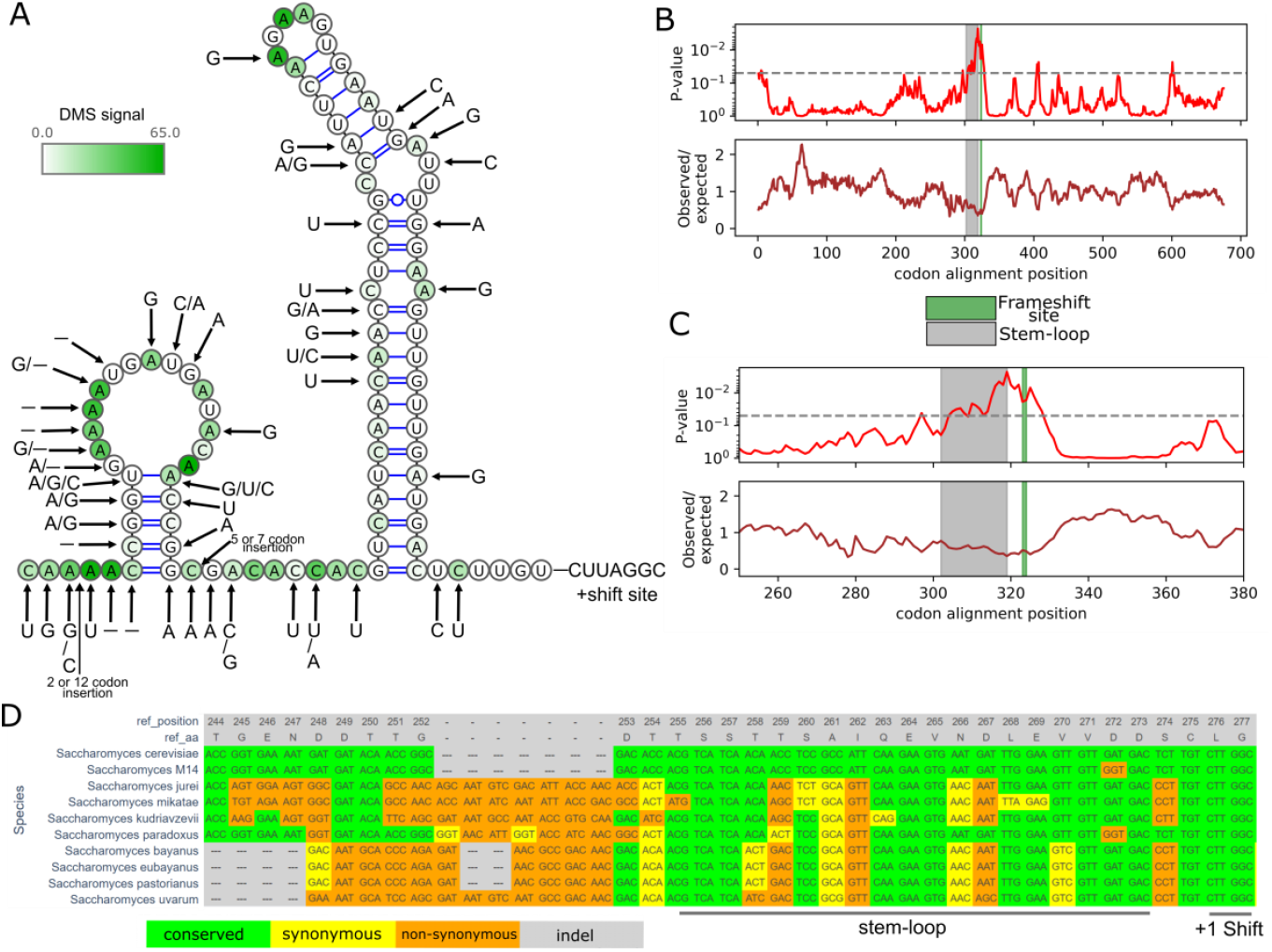
Conserved RNA stem-loop in *ABP140* upstream of the frameshifting site. **A**. RNA secondary structure diagram displaying DMS-seq (chemical targeting of unpaired A and C nts) data supporting the *ABP140* stem-loop. Base substitutions and indels from the *Saccharomyces* are denoted with arrows. **B**. Analysis of the rate of synonymous substitutions with Synplot2 for the ABP140 protein coding region. Observed/expected ratio of substitutions is shown at the bottom track with corresponding p-values above. Dotted line represents p < 0.05 threshold. Below is a zoomed region for the frameshifting cassette. Synplot2 data are visualized from a 17-codon sliding window. **C**. Same as B, except the region is zoomed at the frameshifting site. **D**. Codon alignment of the ABP140 coding sequence with *S. cerevisiae* as a reference.

There is a possibility that *ABP140* frameshifting is stimulated by the nascent peptide within the ribosome peptide tunnel, as discovered in fungal antizyme sequences (48). To test this possibility, we generated mutations (synonymous mutant) that did not change the sequence of the encoded protein but altered the mRNA structure (Fig. 5A). This removed the stimulatory effect suggesting that the stimulation occurs by the RNA structure and not the encoded peptide sequence. Replacing the *ABP140* stem-loop structure with a shorter variant found in *Kluyveromyces marxianus* ortholog (Supplementary Fig. 2) reduces frameshifting to below 50%, but it remains well above 40% in the absence of a stem-loop structure at this position (Fig. 5B). To further exclude the possibility that the upstream sequence induces frameshifting by means other than the described RNA stem-loop structure, we introduced an unnatural stem-loop structure upstream of +1 frameshifting site whose nucleotide sequence does not share sequence similarity with *ABP140* mRNA sequence. This stem-loop structure appears to provide a stimulatory effect similar to wild-type levels (∼55%) (Fig 5B). Progressive truncations from the 5’ end of the *S. cerevisiae ABP140* test sequence also showed that disruption of the stem-loop reduces frameshifting efficiency, and the entire structure is needed for the stimulation (Fig. 5C). In the wild-type mRNA, the base of the stem-loop occurs 2 codons upstream of the frameshifting site. To investigate the importance of the spacer’s length between the stimulatory structure and the frameshifting site, we progressively extended it by up to five codons (Fig. 5D). Increasing the spacer length by just a single codon eliminates the frameshifting-stimulatory effect (Fig. 5D). Collectively, these data provide strong support for the proposition of the RNA structure as frameshifting stimulator and that this is located 5’ of the frameshifting site.

**Figure 5.**
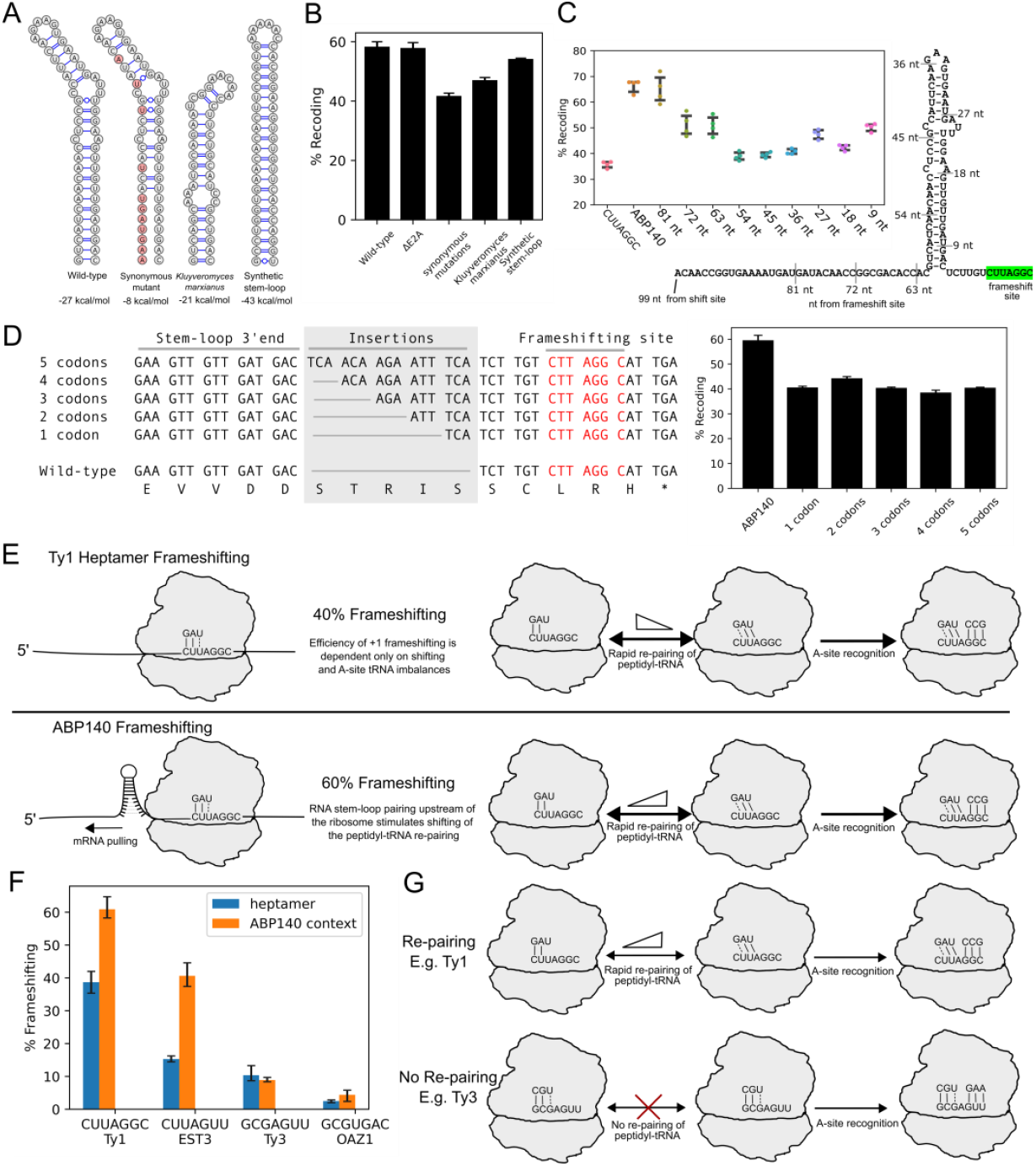
The stimulatory mode of upstream RNA structure affirms two distinct mechanisms of +1 ribosomal frameshifting. **A**. Tested RNA structures, WT, mutants (via synonymous codon substitutions), the stem-loop of *Kluyveromyces marxianus* and the synthetic stem-loop. **B**. Luciferase results showing frameshifting on the *ABP140* test sequence for wildtype and mutants. E2A represents deletion of the second E2A sequence positioned between the test sequence and firefly luciferase coding sequence. **C**. Luciferase assay results showing truncations of the 5’ mRNA context of the *ABP140* test sequence. As a control, the Ty1 heptamer and the original *ABP140* test sequence was tested alongside the truncations. **D**. Schematic of reporters with codon insertions (1-5 codons) between the *ABP140* stem-loop and the frameshift site (left). Recoding efficiencies of these reporters (right). **E**. The proposed model of +1 ribosomal frameshifting by the upstream RNA structure stimulator. **F**. Luciferase assay results showing frameshifting efficiencies of the wildtype heptamers (blue bars) and the heptamers in the context of *ABP140* (orange bars). G. Two models of +1 frameshifting, based on the ability of the P-site to re-pair during +1 frameshifting.

In theory, in addition to frameshifting, the second luciferase could also be expressed via ribosome reinitiation downstream of the first ORF by ribosomes that did not change frame at the frameshifting site. To exclude this possibility, we removed the second StopGo (E2A) site in the vector (between test sequence and Firefly) and carried out Western Blotting analysis of yeast cells expressing this reporter. It showed only a single Firefly luciferase product and no shorter products that would be expected in case of reinitiation (Supplementary Fig. 3). Moreover, the frameshifting efficiency measured with luciferase assays remained the same (Fig. 5B).

### The stimulatory RNA structure promotes P-site tRNA slippage and affirms existence of two distinct mechanisms of +1 ribosomal frameshifting

Most RNA structures stimulating ribosomal frameshifting have been found downstream of frameshifting sites promoting -1 ribosomal frameshifting (4,49,50). In SARS-CoV-2 and likely other instances of ribosomal frameshifting, the stimulation is achieved as a result of the tensions in mRNA between the decoding centre of the ribosome (A and P-sites) and mRNA tunnel entrance where the stimulatory structure is located. This tension prevents the movement of mRNA triplet together with tRNAs during the translocation (51). However, in bacteria frameshifting is often stimulated by Shine-Dalgarno (SD) and anti-SD interactions likely based on a similar principle of tensions within mRNA caused by these interactions. A short distance between the SD and decoding centre pulls mRNA out of the decoding centre promoting +1 frameshifting while a long distance pushes mRNA towards the decoding centre promoting −1 frameshifting (52,53). Furthermore, a somewhat similar phenomenon occurs during transcription in bacteria. Transcription terminators consist of RNA secondary structures upstream of the polyU sequence in the RNA:DNA hybrid within the RNA polymerase bubble and termination involves pulling polyU towards the structure during its formation (54). Transcriptional slippage occurring due to the realignment of RNA relative to its DNA template is also known to be stimulated by upstream RNA secondary structures (55). Inspired by these examples, we proposed a model of how the RNA stem-loop stimulator discovered in this study may stimulate +1 frameshifting during decoding of the *ABP140* mRNA (Fig. 5E). When mRNA is exiting the ribosome, it starts forming the RNA secondary structure eventually pulling mRNA out of the ribosome (in the 5’ direction), thus promoting forward movement of the P-site tRNA and its realignment from the zero frame CUU codon with the +1 UUA codon. In this model, the exact position of the stem-loop would be important and this is supported by the reporters whereby the distance between the stem-loop and frameshifting site is increased (Fig. 5D).

If this model is correct, the RNA secondary structure should only promote +1 frameshifting that involves P-site tRNA movement relative to mRNA (top two mechanisms in Figure 1, but not the bottom one). Thus, we explored the *ABP140* frameshifting stimulator within the context of other known frameshifting heptamers. To achieve this, we substituted the native CUU_A.GG_C frameshifting site with the frameshifting heptamers from *EST3, Ty3*, and *OAZ1*. If the model is correct and we observe a significant increase in frameshifting efficiency, this would imply that all frameshifting sites involve the re-pairing of peptidyl-tRNA to the overlapping +1 frame codon. We observed a substantial increase in +1 frameshifting from 15% to 40% for the EST3 heptamer CUU_A.GU_U, but no significant increase for the Ty3 (GCG_A.GU_U) and OAZ1 (GCG_U.GA_C) heptamers (Fig. 5F). Therefore, these results strongly suggest that ribosomal frameshifting on heptamers with GCG in the P-site do not involve tRNA re-pairing with the +1 codon, and instead, a distinct +1 frameshifting mechanism takes place (Fig. 5G), as originally proposed by Pande *et al* (22).

## DISCUSSION

Prior work on ribosomal frameshifting in *S. cerevisiae* provided widely varying estimates of frameshifting efficiencies, likely because of high variability in assays and the frameshifting cassettes used (56–59). Therefore, to be able to compare different cases of frameshifting, it is important to measure them uniformly. In this work, we set out to explore all known instances of +1 ribosomal frameshifting in *S. cerevisiae* using a combination of approaches: reporter assays, ribosome profiling and comparative sequence analysis. While the aim of the study was to provide a comprehensive and uniform characterisation of frameshifting cassettes across different *S. cerevisiae* genes utilizing ribosomal frameshifting in their expression, several unexpected findings arose.

First, we found a gene, *YJR112W-A*, that is misannotated in the *S. cerevisiae* genome annotation as an intron-containing gene, while it is apparently transcribed into an intron-less mRNA expressing a protein encoded in two ORFs, that are decoded as a single protein via +1 ribosomal frameshifting. This finding illustrates that even nowadays, the protein coding catalogues of eukaryotic genes are not complete even in species with comparatively low frequencies of splicing such as *S. cerevisiae*. Our findings reveal a critical shortcoming in current annotation pipelines: they fail to capture complex mRNA decoding events, largely because the current models assume that all translated regions comprise of a single ORF (60). This finding also demonstrates the utility of ribosome profiling data at overcoming these limitations in identifying novel translated regions.

Second, we unexpectedly identified a novel mechanism of ribosomal frameshifting stimulation. Hitherto frameshifting stimulatory structures were identified downstream of frameshifting sites and are believed to act upon elongating ribosomes by slowing down their movement because of the requirement to unwind these structures (51). Here, we identified a stimulatory structure upstream of the frameshifting site in *ABP140*, suggesting that it acts on the downstream ribosome by promoting its forward movement. Corroborating evidence for this possibility has been provided by a previous study, in which a stem-loop was shown to stimulate +1 frameshifting when artificially placed upstream of the Ty1 frameshifting heptamer in *S. cerevisiae* (61). It is likely that such a mode of frameshifting stimulation is not limited to *S. cerevisiae* and may work in other organisms. Several lines of evidence supporting this come from bacteria. Stimulatory Shine-Dalgarno forming base pairing with rRNA is required for highly efficient +1 ribosomal frameshifting in bacterial release factor 2 (62). Shine-Dalgarno is located at an unusually short distance to the decoding centre and the formed structure likely clashes with the E-site potentially displacing the E-site tRNA during frameshifting (63). Artificial stem-loop structures placed upstream of +1 frameshifting sites stimulate its efficiency *in vitro* (64). Also, a natural upstream secondary structure formed by mRNA exiting the ribosome plays an important role during translational bypassing in bacteriophage gene 60 (65).

Third, we utilized this novel stimulator to address a long-standing question in the ribosomal frameshifting field. How can +1 frameshifting work in the absence of apparent re-pairing of the P-site tRNA with an overlapping P-site codon? Placing the structure upstream of different frameshifting sites revealed its specificity to those frameshifting sites that allow for P-site tRNA re-pairing. In contrast it provided no stimulation at frameshifting sites where re-pairing appears impossible, suggesting that in these cases, frameshifting does not involve repositioning of the P-site codon from the zero to the +1 frame. This finding contradicts a unified model of ribosomal frameshifting in which the forward movement of the P-site tRNA is required regardless of its ability to re-pair with the +1 codon (12). On the contrary, it suggests that a distinct mechanism exists enabling A-site tRNA acceptance at the +1 codon that does not require P-site tRNA repositioning as previously suggested (22). The existence of two such mechanisms may not be specific to yeast, frameshifting sites with seemingly impossible P-site codon re-pairing have been reported in glass sponge mitochondria (66) and under severe limitations of specific amino acids in *Escherichia coli* (67,68).

Finally, while in this study, we focused on the *ABP140* stem-loop, other cases of frameshifting also showed intriguing results. Both the *Ty1* and *EST3* test sequences with native contexts showed a decrease in frameshifting efficiency suggesting that these contexts may have attenuator elements. Surprisingly, *EST3* supposedly contains a stimulatory element (59) surrounding the frameshifting heptamer and we do not observe any such effect with both experimental reporters and publicly available ribosome profiling data analysis.

## Data availability

The data underlying this article are available in the article and in its online supplementary material. Source data, such as reporter readouts of individual experiments are available upon request.

## Acknowledgements

We thank Yousuf Khan (Stanford University) and Sinéad O’Loughlin (UCC) for technical advice in the early stages of this project. We thank Javier Valera (UCC) for supplying the pGREG-505 plasmid.

## Author Contributions

D.A.F. and M.B. carried out experiments. D.A.F. and M.I.S. carried out bioinformatic analyses. D.A.F., J.F.A. and P.V.B. conceived the study. D.A.F., G.L. and M.M.Y. designed experiments and analysed results. D.A.F., J.F.A. and P.V.B. drafted the manuscript and all authors participating in manuscript editing. P.V.B., J.K. and J.F.A. secured funding and provided supervision.

