## Supplementary Fig. for "Comprehensive analysis of yeast +1 ribosomal frameshifting unveils a novel stimulator supporting two distinct frameshifting mechanisms"

**Table of Content:**

Supplementary Figure 1. 2
Supplementary Figure 2 3
Supplementary Figure 3 4

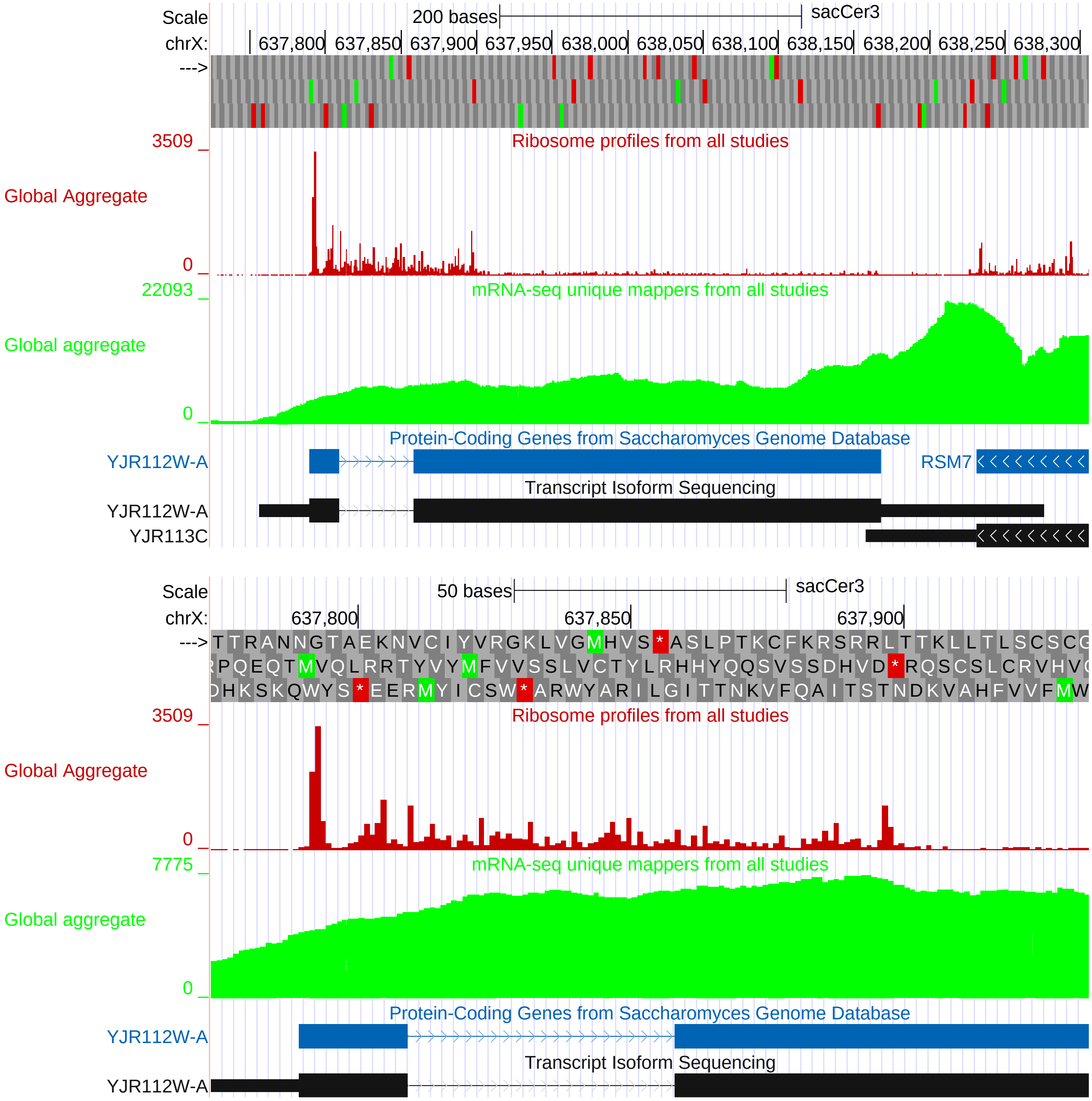


**Supplementary Figure 1.** GWIPS-Viz snapshot of the *S. cerevisiae* YJR112W-A locus (Top). Global aggregate track of Ribo-seq data is presented as red bars reflecting A-site positioning of the translating ribosomes. RNA-seq coverage is presented as the lime (green) track. Bottom panel is the same as top, but with a zoom and focus on the 0-frame ORF that contains the +1 frameshifting sequence CUU_A.GG_C. Note the incorrect annotation tracks that suggest an intronic sequence in the 0-frame ORF.

*Kluyveromyces marxianus*

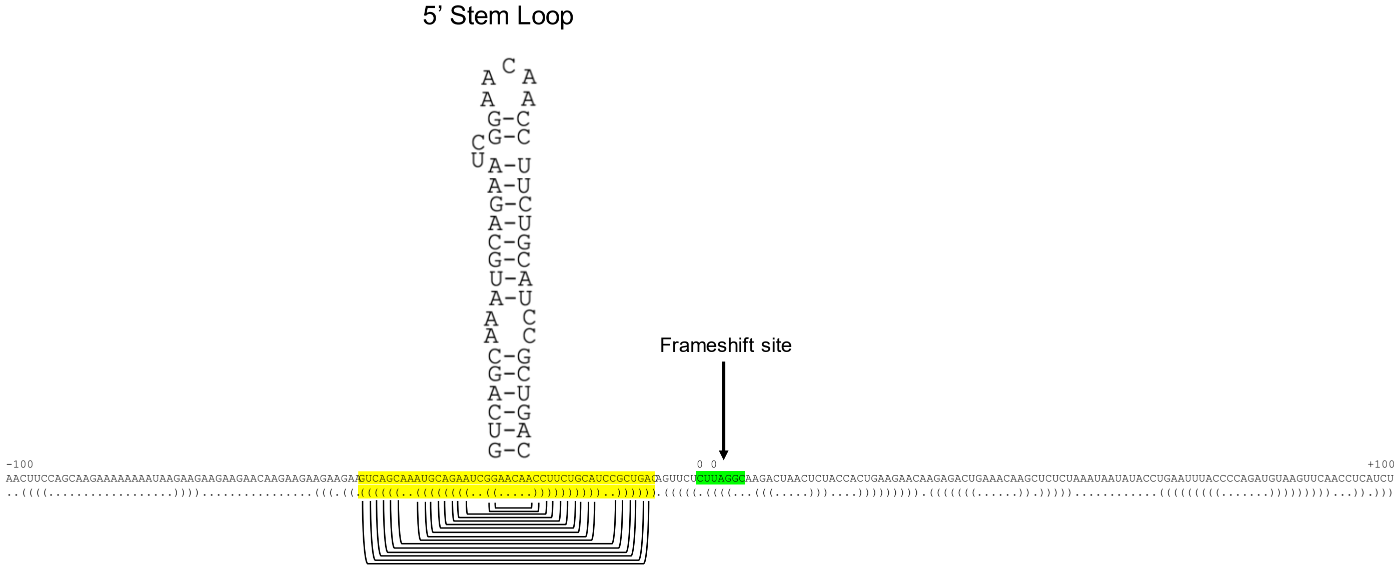


**Supplementary Figure 2**. Sequence of the *Kluyveromyces marxianus* ABP140 mRNA including frameshiftong heptamer (in green) and the 100 nt upstream and downstream. The stem loop located 6 nt upstream from the frameshift site is highlighted in yellow. The predicted secondary structure is indicated below in dot and bracket format, a diagram of of the stimulator’s secondary structure is also shown above.

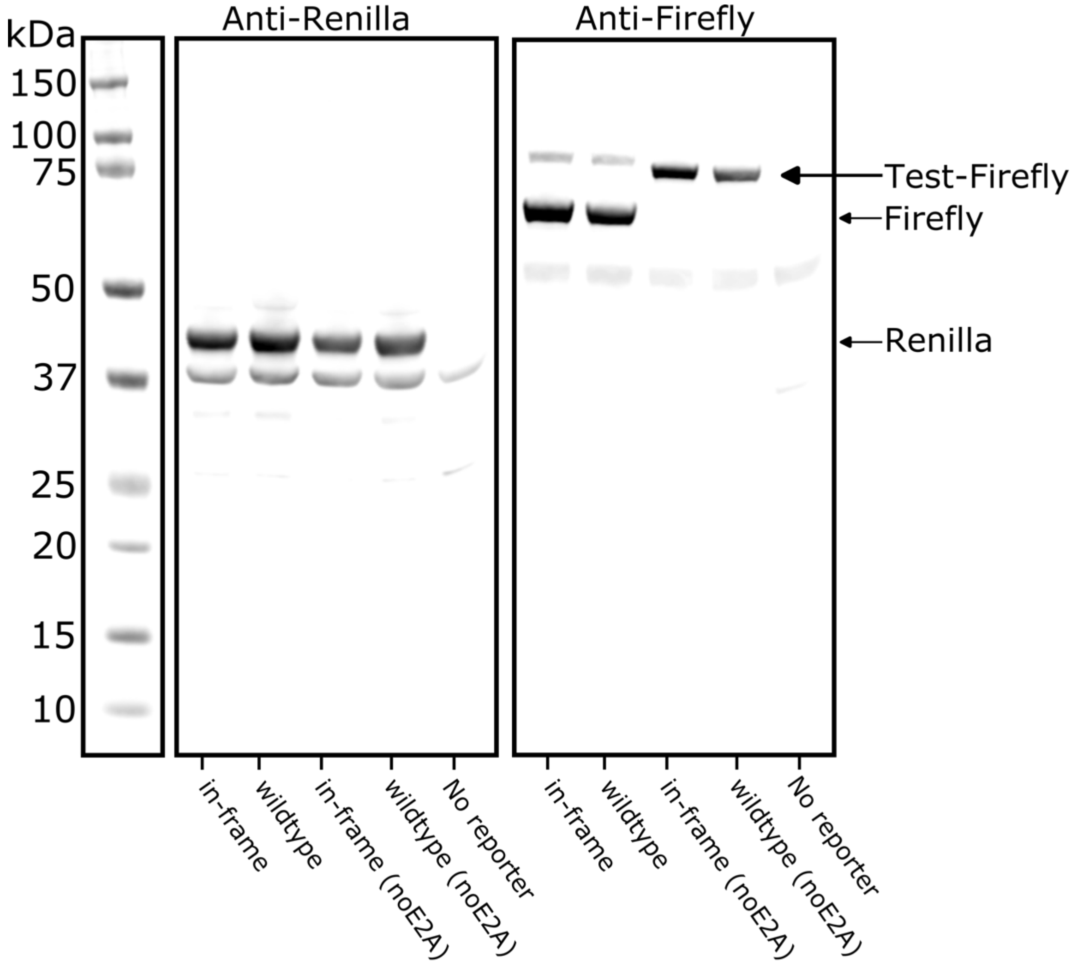


**Supplementary Figure 3**. Western blotting of yeast protein extracts from the cells expressing reporter mRNAs containing ABP140 ribosomal frameshifting cassette and controls. Western blotting was performed with anti-Renilla and anti-Firefly antibodies. As a negative control, yeast cells expressing no plasmid was used. Note that in the noE2A reporter, the downstream (or second) StopGo sequence has been deleted. “Test-Firefly” bands represent the protein products when the second E2A is removed.
